# Across two continents: the genomic basis of environmental adaptation in house mice (*Mus musculus domesticus*) from the Americas

**DOI:** 10.1101/2023.10.30.564674

**Authors:** Yocelyn T. Gutiérrez-Guerrero, Megan Phifer-Rixey, Michael W. Nachman

## Abstract

Parallel clines across environmental gradients can be strong evidence of adaptation. House mice (*Mus musculus domesticus*) were introduced to the Americas by European colonizers and are now widely distributed from Tierra del Fuego to Alaska. Multiple aspects of climate, such as temperature, vary predictably across latitude in the Americas. Past studies of North American populations across latitudinal gradients provided evidence of environmental adaptation in traits related to body size, metabolism, and behavior and identified candidate genes using selection scans. Here, we investigate genomic signals of environmental adaptation on a second continent, South America, and ask whether there is evidence of parallel adaptation across multiple latitudinal transects in the Americas. We first identified loci across the genome showing signatures of selection related to climatic variation in mice sampled across a latitudinal transect in South America, accounting for neutral population structure. Consistent with previous results, most candidate SNPs were in regulatory regions. Genes containing the most extreme outliers relate to traits such as body weight or size, metabolism, immunity, fat, and development or function of the eye as well as traits associated with the cardiovascular and renal systems. We then combined these results with published results from two transects in North America. While most candidate genes were unique to individual transects, we found significant overlap among candidate genes identified independently in the three transects, providing strong evidence of parallel adaptation and identifying genes that likely underlie recent environmental adaptation in house mice across North and South America.

**Author summary:** Since their arrival with European colonizers, house mice have successfully spread throughout the Americas. There is strong evidence that populations in North America have adapted in that time, including parallel evolution of phenotypes across latitude (e.g., body size, behavior) as well as the identification of genes that show signals of selection. Here, we investigate the genetics of environmental adaptation in South America. We find that populations in South America evolve independently of populations in North America. We identify candidate genes for environmental adaptation with links to traits like body size, metabolism, immunity, eye function, and the cardiovascular and renal systems. We then bring together data from three transects across two continents to determine if environmental adaptation is predictable, with parallel genetic changes in response to shared conditions. We find that most evidence of environmental adaptation lies in regulatory regions and that, while most candidate genes are unique to individual transects, many are shared, providing significant evidence of parallel adaptation. We identify a core set of candidate genes independently identified in all three transects that likely contribute to environmental adaptation in the Americas. These results highlight the value of studying wild populations of this genetic model system.

## Introduction

Understanding the genetic details of how species adapt to new environments is a key goal of evolutionary biology. One approach to investigating the genetic basis of environmental adaptation is to look for covariation between allele frequencies and environmental variables (Huxley 1938; Endler 1977). Such clines can result from neutral processes, but statistical methods can be used to account for neutral population structure (e.g., Coop *et al*. 2010; Frichot *et al*. 2013) and this approach has been applied successfully to a wide range of organisms (e.g., Hancock *et al*. 2010; Fournier-Level *et al*. 2011; Bohutínská *et al*. 2021; Magalhaes *et al*. 2021; Chaturvedi *et al*. 2022). An extension of this approach is to compare patterns of genetic variation across multiple independent environmental gradients (e.g., Umina *et al*. 2005; Hohenlohe *et al*. 2010; Paaby *et al*. 2010; Adrion *et al*. 2015; van Boheemen and Hodgins 2020; Chaturvedi *et al*. 2022). For example, comparisons of *Drosophila melanogaster* populations in the northern and southern hemispheres have identified shared responses to selection (Reinhardt *et al*. 2014; Adrion *et al*. 2015; Juneja *et al*. 2016). When neutral population structure is accounted for, parallel clines can provide strong evidence that particular genes and traits contribute to adaptation even when the specific mechanism is unknown. Characterizing phenotypic variation in wild populations can be difficult, and biologically important phenotypes contributing to adaptation may go undetected. Even when there are known clines in phenotypes, many of the traits of interest may be polygenic and influenced by the environment. Detecting signatures of selection on complex traits and connecting those changes to phenotypes remains challenging (Hermisson and Pennings 2005; Messer and Petrov 2013; Berg and Coop 2015; Harris *et al*. 2018; Barghi *et al*. 2020). However, parallel clines can help identify particular genes and traits that are under selection. Moreover, because genome scans are agnostic with respect to phenotype, this kind of comparative approach can also point to previously unnoticed traits that may be important to adaptation.

House mice (*Mus musculus domesticus*) provide an opportunity to study the genomic basis of environmental adaptation using natural replicates. Native to Western Europe, house mice have spread opportunistically around the world in association with humans during the last five hundred years (Boursot *et al*. 1993; Berry *et al*. 1978; Hardouin 2010; Morgan 2022; Agwamba and Nachman 2023). In this short time, they have successfully colonized both North and South America, from Tierra del Fuego (55°S) to Alaska (61°N), spanning an enormous range of habitats and climates. Previous studies have found that house mice exhibit clinal variation in body size, with size increasing with distance from the equator in South America and North America, consistent with Bergmann’s Rule (Lynch 1992; Phifer-Rixey *et al*. 2018; Suzuki *et al*. 2020, Ferris et al. 2021; Ballinger and Nachman 2022). House mice also show clines in ear length and tail length across North and South America, with length decreasing with increasing distance from the equator, consistent with Allen’s Rule (Ballinger and Nachman 2022). These observations conform to well-known ecogeographic patterns in mammals and are thought to reflect thermoregulatory adaptations for animals living in cold or warm environments. These differences persist in a common laboratory environment for multiple generations, indicating that they are genetically based (Phifer-Rixey *et al*. 2018; Ferris et al. 2021; Ballinger and Nachman 2022). Genomic surveys have identified candidate genes using covariation between environmental variables and genetic variation in two clines across latitude in North America (Phifer-Rixey *et al*. 2018; Mack *et al*. 2018; Ferris *et al*. 2021) in tandem with phenotype and gene expression data (Phifer-Rixey *et al*. 2018; Mack *et al*. 2018). Furthermore, Ferris *et al*. (2021) compared results from the two latitudinal transects in North America and identified significant overlap in signals of selection, including at several genes related to heat sensing (*Trpm2*) and body weight (*Mc3r* and *Mtx3*), suggesting some shared response to selection.

Less is known about genetic variation in South American populations. Previous studies of altitudinal adaptation (Storz *et al*. 2007; Beckman *et al*. 2022) and cytogenetics (Giménez and Bidau 1994) provided evidence that house mice in South America derive from the same subspecies (*M. m. domesticus*) as mice in North America. However, sampling was limited, and it is not known whether there may be introgression from other subspecies. Patterns of genetic diversity and differentiation across the continent are also largely unknown as are the relationships to populations in North America. Importantly, some aspects of the environment, such as temperature, vary similarly across latitude in North and South America (Paruelo *et al*. 1995) providing an opportunity for an investigation of parallel rapid environmental adaptation across two continents.

Here, we explore genomic signatures of environmental adaptation in house mice from South America across a latitudinal transect from equatorial Brazil (∼3° S) to southern Argentina (∼ 55° S) using exome capture data of wild-caught individuals. We combined the data generated in this study with published data from eastern and western North America. We address five main questions. First, do house mice in South America derive from the same subspecies (*M. m. domesticus*) as house mice in North America? House mice comprise three major subspecies which diverged ∼150,000 – 500,000 years ago and have distinct ranges: *M. m. musculus* is found in Eastern Europe and northern Asia, *M. m. domesticus* is found in Western Europe and the Mediterranean region, and *M. m. castaneus* is found in SE Asia (e.g., Suzuki 2004, 2013; Geraldes *et al*. 2008, 2011; Duvaux *et al*. 2011; Phifer-Rixey *et al*. 2020). *M. m. domesticus* is the presumed source population for the Americas (Boursot *et al*. 1996; Gabriel *et al*. 2010; Suzuki *et al*. 2013, Didion and Pardo-Manuel de Villena 2013), although the subspecific origin of house mice across most of South America has never been explored. Second, do house mice in North America and South America have independent evolutionary histories? If so, this would suggest that any shared signals of selection result from parallel evolution. Third, which genes show signatures of selection in house mice from South America? Fourth, to what extent are parallel signatures of selection seen in comparisons among mice from three different latitudinal transects: South America (SA), eastern North America (ENA), and western North America (WNA)? Previous work showed that mice in eastern and western North America have independent evolutionary histories (Ferris et al. 2021), providing an opportunity here to compare three phylogenetically independent transects. Finally, what genes underlie variation in body size and are they associated with signals of selection? We found that mice in South America are of *M. m. domesticus* origin and that they have an independent evolutionary history from mice in North America. We also found signatures of selection across the genome among mice from South America and significant overlap among candidate genes for all three transects, providing evidence of parallel adaptation. Finally, a genome-wide association study (GWAS) identified eight genes associated with differences in body weight, all except one of which showed signatures of selection.

## Results

### *Mus musculus domesticus* ancestry in the Americas

We sequenced the complete exomes of 86 wild house mice sampled from 10 populations along a latitudinal transect from central Mexico to the southern tip of South America (Fig. 1a, Table S1). To analyze patterns of admixture, we combined these data with previously published data from populations in eastern (n=50) and western (n=50) North America (Phifer-Rixey *et al*. 2018; Ferris *et al*. 2021) and published data from each of the three major *Mus musculus* subspecies (Table S2; Harr *et al*. 2016). Specifying K=3 genetic clusters, we found that house mice in the sampled populations of the Americas are of *M. m. domesticus* origin, apart from one population in Tucson which is mostly of *M. m. domesticus* origin but also shows some limited admixture with *M. m. castaneu*s, as previously reported (Fig. 1b; Ferris *et al*. 2021). These results provide strong evidence that house mice in Mexico and South America are *M. m. domesticus* with no evidence of significant introgression from the other subspecies.

**Figure 1.**
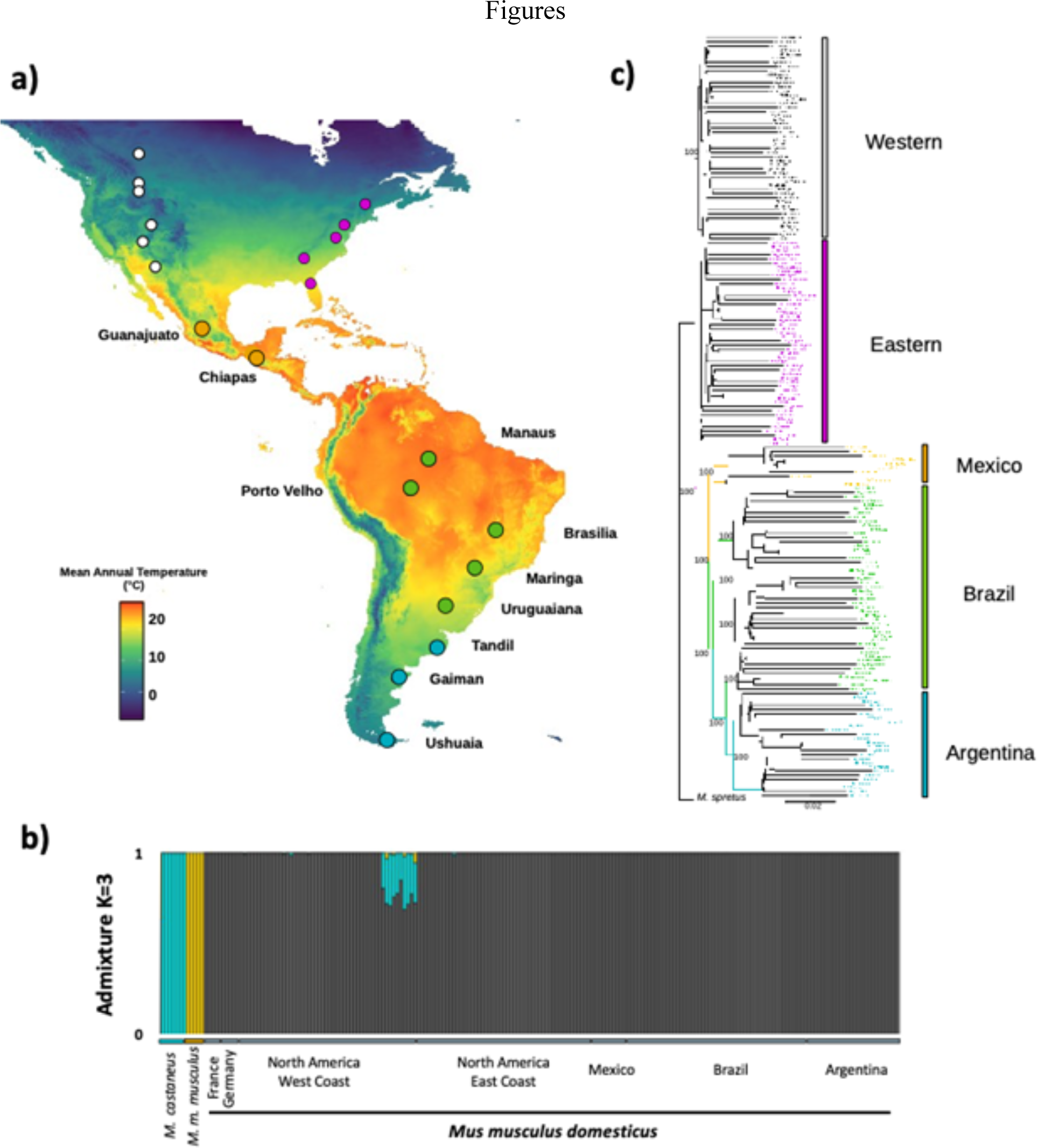
**a**) Map of mean annual temperature across the Americas (WorldClim database). Populations of wild house mice sampled across a latitudinal transect in Mexico and South America localities are shown with large circles. Populations included in previously published surveys in North America (Phifer-Rixey *et al*. 2018; Ferris *et al*. 2021) are shown with small circles. **b)** Admixture plot including representatives from all three primary subspecies of house mouse as well as mice from sampled populations in the Americas. **c**) Phylogenetic reconstruction of *Mus musculus domesticus* populations across the Americas, with *M. spretus* as the outgroup.

### Independent recent evolutionary history of sampled transects

We constructed a maximum likelihood phylogenetic tree using RAxML (Stamatakis 2014) with *M. spretus* as an outgroup (Fig. 1c, Table S2). For this analysis, we pruned the dataset to only include autosomal sites for which 80% of the individuals were covered, resulting in 895,333 sites. This analysis identified three major clades: populations from western North America, populations from eastern North America, and populations from Mexico and South America, each with 100% bootstrap support (Fig. 1c). Within South America, mice formed two reciprocally monophyletic groups, each with 100% bootstrap support, largely corresponding to a northern clade (Manaus, Porto Velho, Brasilia, and Maringa) and a southern clade (Uruguaina,Tandil, Gaiman, and Ushuaia). In North America, mice formed two reciprocally monophyletic groups, each with 100% bootstrap support, corresponding to the eastern and western transects as previously reported (Ferris *et al*. 2021). Thus, these analyses suggest that mice in each of the three transects have an independent evolutionary history.

### Population structure in the Americas

We first used NGSadmix to explore patterns of population structure within South America (Fig. 2a), excluding Mexico because of limited sampling. The number of separate clusters that best fit the data was 5 (Evanno test). At K=5 we observed clusters that corresponded to each sampled population or geographically close pairs of populations (Manaus and Porto Velho, Brasilia and Maringa, and Tandil and Ushuaia). At K=8, each sampled population formed its own cluster. Consistent with the phylogenetic tree, principal component analysis including all populations in the Americas revealed genetic differentiation between North and South America, in which PC1 and PC2 largely separate populations by latitude (Figs. 1c, 2b). When the first three principal components are plotted together, five major clusters are observed, corresponding to eastern North America, western North America, Mexico, northern South America, and southern South America.

**Figure 2.**
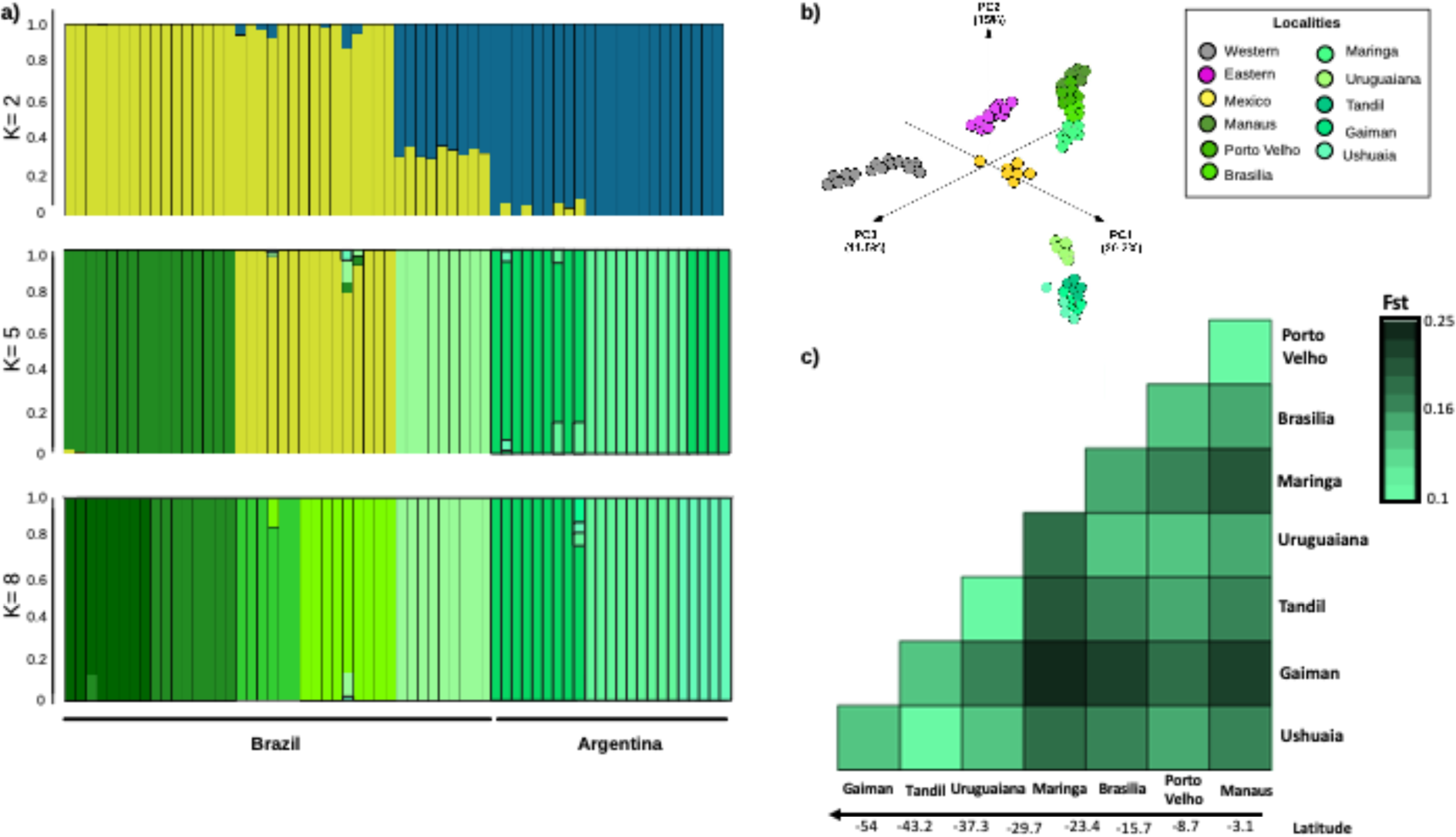
**a)** Admixture plot from South American populations evaluating K=2, 5, and 8. **b)** Genetic PCA of *Mus musculus domesticus* populations across the Americas. **c)** Heatmap of pairwise genetic differentiation (*F_st_*) values between the eight populations in South America.

The average *F_st_* in pairwise comparisons among all eight South American populations (*F_st_* = 0.154; Fig. 2c) was significantly higher than the average pairwise *F_st_* seen among the 10 North American populations (*F_st_* _Eastern_ = 0.069, Mann-Whitney U *p-value* = 0.00001, *z-score* = 4.52, and *F_st_* _Western_ = 0.079, Mann-Whitney U *p-value* = 0.00001, *z-score* = 4.6; Table S3). We also detected a significant signature of isolation by distance among populations in SA (IBD; *Mantel statistic: R^2^* = 0.2246, *p* = 0.0001; Fig. 2c; Table S3) in contrast to only modest evidence of isolation by distance in the WNA transect (Ferris *et al*. 2021) and no evidence in the ENA transect (Phifer-Rixey *et al*. 2018). The distance between sites is larger in the South American transect. However, even when limiting comparisons to similar geographic distances, there is a significant positive correlation between geographic and genetic distance in SA (y = 4×10^-5^x+ 0.091, *R^2^*=0.42), while there is no evidence of a correlation in ENA (y= -3×10^-6^ x +1.37, *R^2^*=0.0005). These differences suggest that barriers to gene flow (physical, political, or otherwise) may differ between the continents.

### Genomic signatures of environmental adaptation in house mice from South America

We first explored variation in climatic variables across the sampled localities in South and North America. In a PCA analysis of the 19 bioclimatic variables from the WorldClim database, the first two principal components explained 71.53% of the total variance (Fig. S1a; Tables S4-S5). As expected for latitudinal sampling in this region, the first principal component was largely driven by mean annual temperature (MAT, Bio1) and other highly correlated temperature related variables. The second principal component was mainly associated with precipitation of the driest month (PDM, Bio14) and precipitation of the driest quarter (Bio17; Fig. S1a). Since our sampling design was based on a latitudinal transect, we were most interested in exploring the effects of climatic variables associated with PC1, for which MAT showed a significant negative correlation with latitude (*R^2^*=0.85, *p-value* ≤ 2.2 x 10 ^-16^; Fig. S1b). Mean annual temperature showed a very high degree of overlap, varying from near 5°C to over 20°C in all three transects. In contrast, precipitation of the driest month showed variation in all three transects, but little overlap among transects (Figs. S1c, S1d).

To identify candidate SNPs that show signals of environmental adaptation in South America, we used a latent factor mixed model (LFMM) that implements a Bayesian bootstrap algorithm and accounts for population structure (Frichot *et al*. 2013). To avoid including close relatives, we estimated the pairwise relatedness coefficient between individuals from the same population using NgsRelate (Hanghøj *et al*. 2019). Individuals with a pairwise relatedness value greater than 0.25 were considered close relatives and were removed from the analysis (Fig. S2). Of the 76 mice from South America, 14 had a pairwise relatedness value greater than 0.25 and were removed. Next, we also excluded individuals from Brasilia and Ushuaia due to small sample sizes (N=5) for these populations, leaving 52 individuals from six populations (Manaus N=8, Porto Velho N=8, Maringa N=9, Uruguaiana N=9, Tandil N=9, and Gaiman N=9). Given those populations, we filtered SNPs retaining those with a minimum allele frequency of 5% and with sites called in at least 80% of all individuals (270,720 SNPs). We then conducted genomic scans for selection with LFMM at K=3, first with latitude and then with MAT and PDM. We identified sites |*z-score|* ≥ 2 and a *q-value ≤* 0.01 after False Discovery Rate (FDR) correction as outliers. Genes that contained outlier SNPs were considered candidate genes in all following analyses.

We identified 9,600 outlier SNPs in >3,400 genes across the genome associated with variation in latitude and 7,481 outlier SNPs in >2,800 genes for MAT (Figure 3b; Tables 1, S6). We identified far fewer candidate SNPs for PDM (1,0007 SNPs in ∼500 genes) which was expected given that the sampling scheme was not designed to explore variation in this variable (Table S7; Figure 3a, b). The vast majority of candidate SNPs were not amino-acid changing (Figure 3a; Table S7) and most genes that contained a candidate SNP did not contain other SNPs that were amino-acid changing (Table S7). These results are consistent with results from the eastern and western transects of North America scans using latitude and mean annual temperature as variables (Phifer-Rixey *et al*. 2018; Ferris *et al*. 2021).

**Figure 3.**
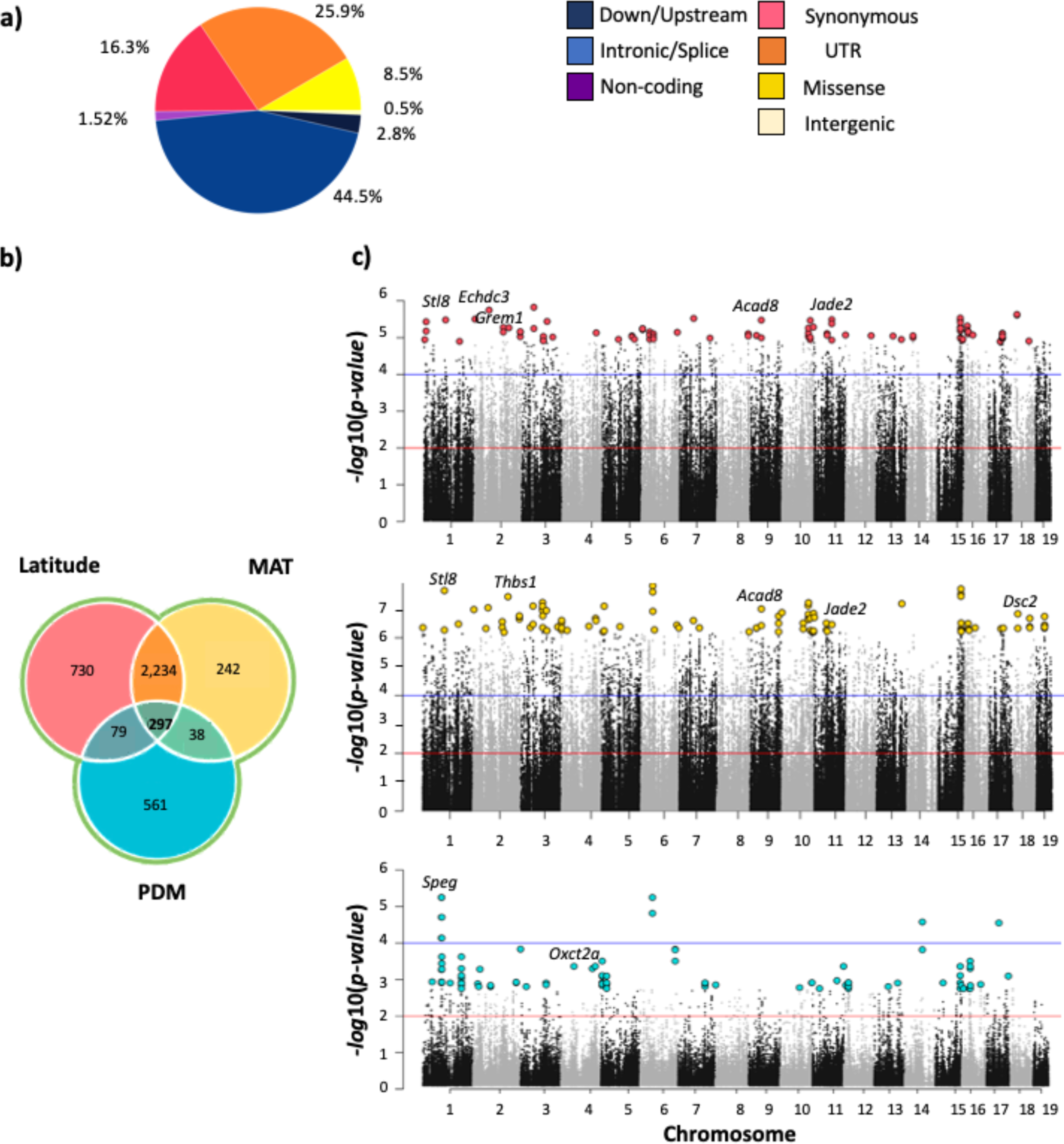
Signatures of selection among house mice from South America. **a)** The distribution of outlier SNPs across predicted variant effect categories. Proportions shown are averages across LAT, MAT, and PDM; full data are given in Table S7. **b)** Venn diagram showing the number of unique and shared candidate genes in the South America transect for latitude, mean annual temperature (MAT), and precipitation of driest month (PDM). **c)** Manhattan plots showing the results of the population genomic scans for selection for the three environmental variables (red line indicates *q-value* = 0.05, and blue line indicates a *q*-*value* = 0.001). Highlighted are the first 100 hundred SNPs with the lowest *p-values* for each environmental variable.

To explore the functional significance of candidates, we conducted gene ontology enrichment analyses for each set of candidate genes (topGO R package; Alexa and Rahnenfuhrer, 2022; Table S8), used MouseMine (MGI) to identify phenotypes associated with the genes (Motenko *et al*. 2015), and Panther to explore pathways (Thomas *et al*. 2022). For latitude, we found significant enrichment (classic Fisher correction *p* ≤ 0.05) in a variety of biological processes including those related to intracellular transport, heart development and function, retina development, lipid metabolism, behavior, and brain development (Table S8). We also identified top candidate genes for latitude (genes annotated to the 100 SNPs with the lowest *p-values*) for which mutants are associated with phenotypes related to body weight and size, fat, metabolism (lipids, cholesterol, insulin, triglycerides, leptin, etc.), immunity, cardiac function, limb and organ morphology, locomotion, and eye function/development, among others (Figure 3c; Tables 1, S6, S8). Functional results for MAT were similar to those for latitude (Figure 3c; Tables 1, S8). For PDM, we detected candidate genes with diverse mutant phenotypes including those related to immunity, muscle function, and kidney function/morphology (Figure 3c; Table S8). There was enrichment of biological process GO terms including those related to sensory perception/response and cholesterol, among others (Figure 3c; Table S8).

As expected, most of the candidate SNPS and genes identified using latitude and MAT were shared (>6,541 shared SNPs from 2,545 genes and 2,575 shared genes including those for which there was not exact overlap at the SNP level), whereas candidates identified using PDM were largely unique (Fig. 3b, c). Candidates shared among latitude and MAT were significantly enriched for many gene ontology annotations related to biological regulation, heart development, eye development, lipid metabolism, and others (Table S8). Twenty-three genes were annotated to the 100 SNPs with lowest corrected p-values for both latitude and MAT (Table 1). These genes are linked to many mammalian phenotypes (MGI, Motenko *et al*. 2015; Tables 1, S8) including abnormal blood homeostasis (*Bcl2l11, Grem1, Jade2, Kif21a, Nlgn2, Plscr3, Sar1b, St18, Tbl1xr1*), lipid and fatty acid metabolism, body fat, and related phenotypes (*Bcl2l1*, *Plscr3, Sar1b, St18, Tblxr1*), abnormal glucose homeostasis *(Car13, Jade2, Kif21a, Plscr3, Tbl1xr1),* immunity (*Arrdc4, Bcl2l11, Cyren, Jade2, St18*), abnormal eye/retina morphology (*Car13,Dsc2, Dscaml1, Kif21, St18)*, abnormal renal/urinary system (*Arrdc4*, *Bcl2l11*, *Cald1, Grem1, Jade2*), abnormal cardiovascular system morphology (*Cyren, Dsc2, St18, Tbl1xr1*), abnormal limb and digit morphology (*Grem1*), and behavior (*Nlgn2*) (Table 1). Four of these 23 genes included top candidate SNPs classified as missense modifications (*Agbl3, Cald1, Vwa5b2, Vps8).* In humans, *Plscr3* and *Sar1b* are linked to obesity and lipid metabolic disease, respectively, *Kif21a* is linked to ocular disease, and *Nlgn2* is linked to schizophrenia (MGI, Motenko *et al*. 2015).

**Table 1.**
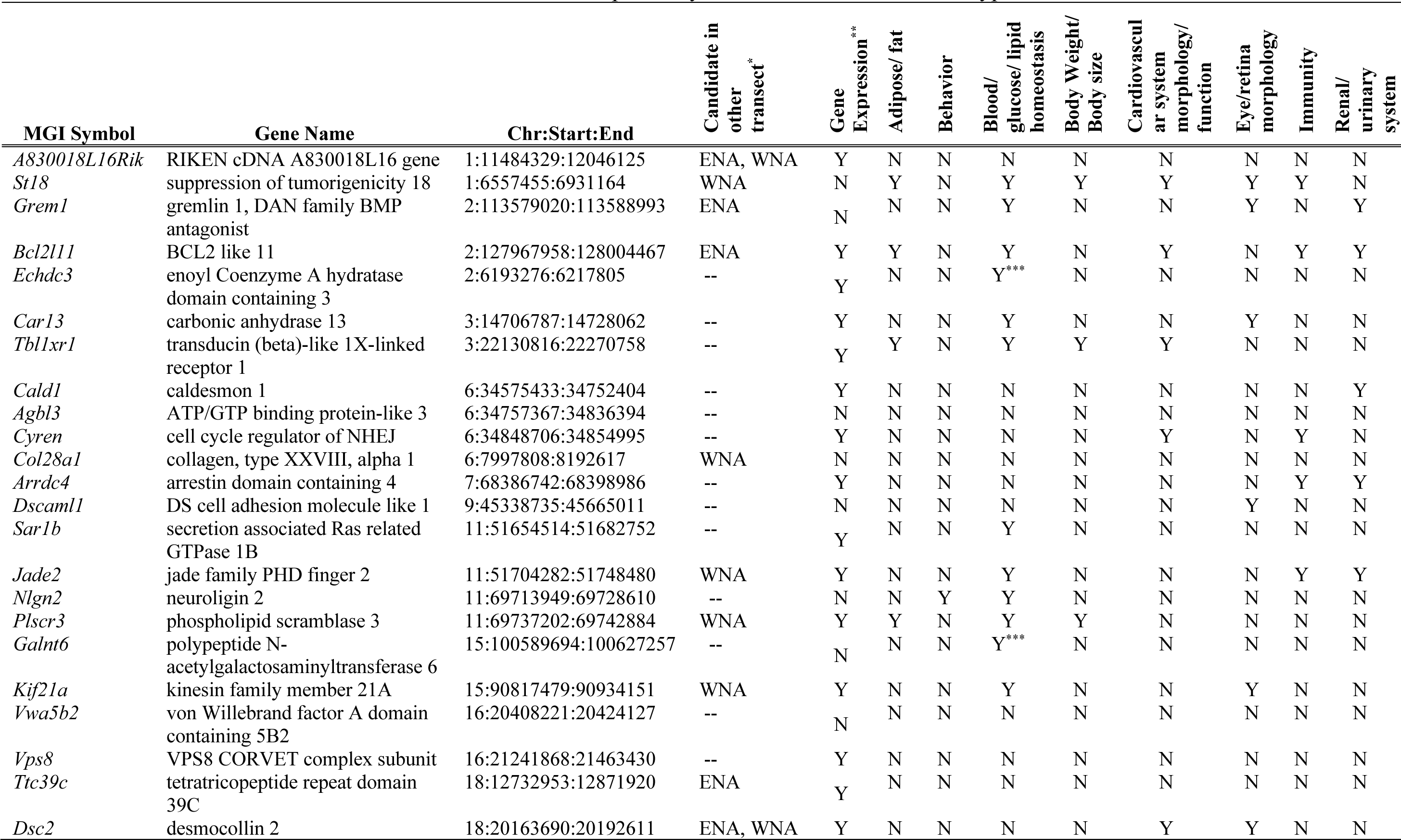
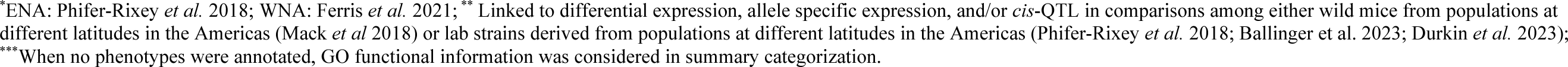
Genes annotated to top candidate SNPs in LFMM analyses of both latitude and mean annual temperature in the South American transect. Functional summarization is based primarily on MGI Mammalian Phenotype annotations and is not exhaustive.

### Parallel adaptation across transects in the Americas

The three transects were planned to capture responses to climatic variables that covary with latitude, allowing us to investigate the extent to which responses to selection are shared. As expected, all three transects have similar ranges in MAT, from near 5°C to over 20°C, although the shift is more gradual in South America, covering ∼50 degrees of latitude compared to ∼20 degrees of latitude in the other transects (Fig. 1a; Table S5). The transects were not planned to cover clines in PDM and there was little overlap in the range of values among transects in PDM (Fig. S1a, c, d), but we also considered this variable given its inclusion in the analysis of South American populations and since it reflects climatic variation that is largely orthogonal to MAT (Fig. S1a) To ensure standardization, we repeated genome scans using published exome-capture data for transects in eastern North America (50 individuals in five populations; Phifer-Rixey *et al*. 2018) and western North America (50 individuals in five populations; Ferris *et al*. 2021; Fig. 1a; Table S1) using exactly the same approach to calling SNPs and genotypes as for the South American transect. After filtering, there were 281,362 SNPs in the transect from eastern North America and 342,108 SNPs in the transect from western North America (minimum allele frequency of 5%, with sites called in at least 80% of all individuals). For each transect and variable (latitude, MAT, and PDM), we identified candidate SNPs applying a *q-value* cut-off ≤ 0.05 and |*z-score| ≥* 2 (Table S9). To identify shared responses to selection among transects, we compared the candidate genes identified using each environmental variable in pairwise comparisons between transects (Table S10). We performed permutation tests using 100,000 replicates with replacement to evaluate whether the overlap in shared genes in pairwise comparisons was significantly greater than expected by chance (Table S11).

In each of these three pairwise comparisons, the number of shared candidate genes was significantly greater than expected by chance in all analyses of latitude and MAT (*p-value ≤* 0.005, *z-score* ≥ 3 for each of the six comparisons; Tables S10, S11). Overlap among candidates identified using PDM was significantly more than expected by chance in only one comparison (WNA-ENA, *p-value =* 0.007, *z-score* = 2.65; Tables S10, S11). The proportion of shared genes was also much higher for latitude (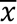 *=* 16.50%) and MAT (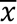*=* 16.22%) than for PDM (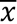*=* 6.95%; Fig. 4a). When considering the overlap among all three transects, the pattern was even more pronounced. The proportions of genes shared among all three transects for latitude (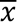= 7.30%) and MAT (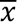*=* 7.16%) was more than seven times greater than for PDM (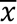*=* 0.93%; Table S10). These observations are consistent with the overlap in climatic variables relating to temperature but not precipitation among transects (Fig. 4a). Nevertheless, while there was more overlap among transects than expected by chance for latitude and MAT, most signals of selection were specific to individual transects (Fig. 4a).

**Figure 4.**
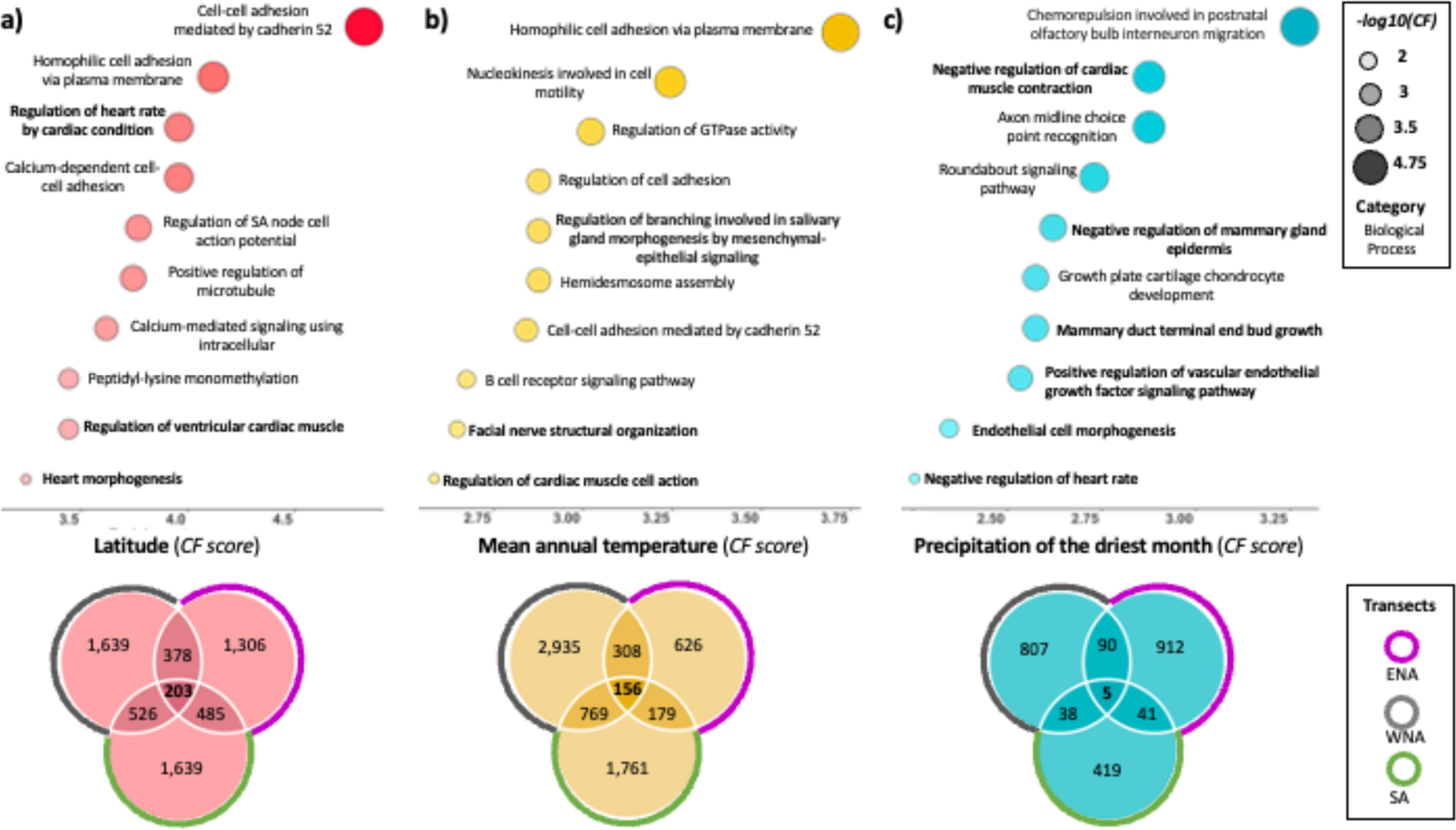
Venn diagram illustrating the shared candidate genes across the three transects [South America (SA), Eastern of North America (ENA), and Western of North America (WNA)], and the significantly enriched gene ontologies for each environmental variable: **a)** latitude, **b)** MAT and **c)** PDM (complete enrichment information is on Supplementary Table S11). CF (*Classic fisher* test used for the enrichment analysis).

Finally, we explored the function of candidate genes shared among the three transects for each environmental variable (e.g., Motenko *et al*. 2015; Thomas *et al*. 2022). For latitude, 203 genes were shared in the three transects with diverse annotated phenotypes (MGI, Motenko *et al*. 2015) including those related to metabolism (insulin, glucose, leptin, lipids, cholesterol, etc.), body size/fat, immunity, reproduction, eye development/function, behavior, and cardiovascular function/development. There was a significant enrichment of gene ontologies associated with regulation of heart rate/cardiac muscle, immunity, the eye, and calcium ion transport, among others (Fig. 4a; Tables S9, S11, S12). Links to cardiovascular function suggest possible impacts on thermoregulation (for example, an efficient mechanism to avoid blood vessel constriction when temperature drops; Withers *et al*., 2016; Tan and Knight 2018). There was also a significant enrichment of genes in Reactome pathways related to collagen formation and degradation, axon guidance, and MET activates PTK2 signaling. For MAT, we found that 156 genes were shared, with a similar range of functions as for latitude (Fig. 4b; Table S10, S11, S12). Only five candidate genes were shared among transects for PDM (*Cgnl1, Col27a1, Myo15, Robo1, Sri*; Figure 4*c*).

In total, 90 genes were identified for both latitude and MAT in all three transects (Table S12). Those 90 genes include many annotated to knock-out/mutant phenotypes for abnormal homeostasis (43), abnormal blood homeostasis (29), abnormal motor/coordination, movement (27), abnormal postnatal growth/weight/body size (23), abnormal cardiovascular morphology (21), abnormal immune system morphology (20), and abnormal glucose homeostasis (17), among others. GO annotations are also diverse, but many have biological processes annotations related to high level terms like regulation of biological processes (62), response to stimulus (51), metabolic processes (48), developmental processes (41), gene expression (21), immune processes (10), and growth (10). Of these 90 genes, 24 were top candidates (annotated to the 200 SNPs with lowest *p*-values) in at least one transect. Many of these have mutant phenotypes related to abnormal behavior (12), abnormal homeostasis (11) and related child terms, abnormal immune system morphology (6), and abnormal body size (6). Over half have GO annotations related to metabolic processes (14). Notable among these are genes like *Mc3r* which was identified as a top candidate in both North American transects and is functionally linked to body size and metabolism (Table 2). *Rorb* is a nuclear receptor that functions in photoreceptors. Normal development of rod and cone photoreceptor cells depends on *Rorb* (e.g., Jia *et al*. 2009) and it is known to be involved in the regulation of circadian rhythm (e.g., André et al. 1998; Masana *et al*. 2007). *Rorb* was a top candidate in two transects and linked to differences in gene expression in strains derived from the Americas (Ballinger *et al*. 2023). *Met* was identified as a top candidate in two transects, has a wide variety of annotated phenotypes including those relating to metabolism, cancer, and pulmonary function/morphology, and showed differential expression among new wild derived strains from different latitudes in the Americas (Ballinger *et al*. 2023; Durkin *et al*. 2023). *Akap9* which was a top candidate in the South American transect, is also linked to differences in gene expression in strains derived from the Americas (Phifer-Rixey *et al*. 2018; Ballinger *et al*. 2023, Durkin *et al*. 2023) and has annotated phenotypes relating to fat, immunity, and metabolism.

**Table 2.**
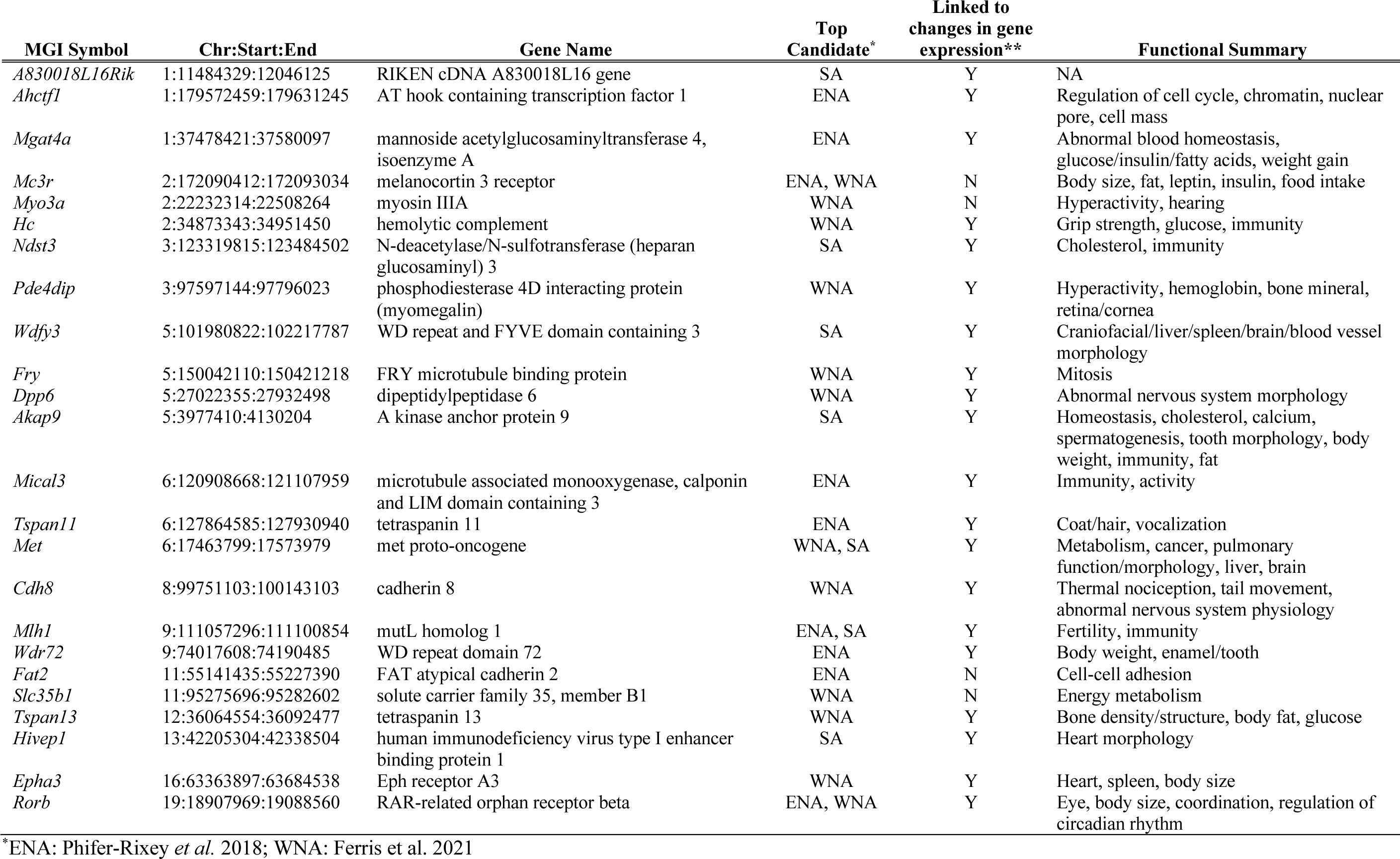
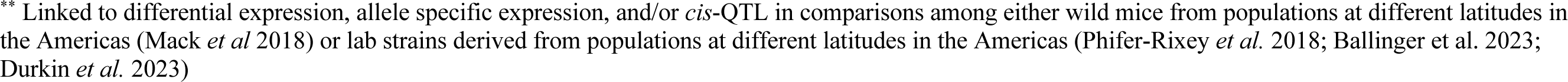
Candidate genes shared among all three transects for both latitude and mean annual temperature that were a top candidate for at least one of the variables in at least one transect. Functional summarization is based primarily on MGI Mammalian Phenotype and GO Biological Process annotations and is not exhaustive.

| MGI Symbol | Chr:Start:End | Gene Name | Top Candidate* | Linked to changes in gene expression** | Functional Summary |
| --- | --- | --- | --- | --- | --- |
| *A830018L16Rik* | 1:11484329:12046125 | RIKEN cDNA A830018L16 gene | SA | Y | NA |
| *Ahctf1* | 1:179572459:179631245 | AT hook containing transcription factor 1 | ENA | Y | Regulation of cell cycle, chromatin, nuclear pore, cell mass |
| *Mgat4a* | 1:37478421:37580097 | mannoside acetylglucosaminyltransferase 4, isoenzyme A | ENA | Y | Abnormal blood homeostasis, glucose/insulin/fatty acids, weight gain |
| *Mc3r* | 2:172090412:172093034 | melanocortin 3 receptor | ENA, WNA | N | Body size, fat, leptin, insulin, food intake |
| *Myo3a* | 2:22232314:22508264 | myosin IIIA | WNA | N | Hyperactivity, hearing |
| *Hc* | 2:34873343:34951450 | hemolytic complement | WNA | Y | Grip strength, glucose, immunity |
| *Ndst3* | 3:123319815:123484502 | N-deacetylase/N-sulfotransferase (heparan glucosaminyl) 3 | SA | Y | Cholesterol, immunity |
| *Pde4dip* | 3:97597144:97796023 | phosphodiesterase 4D interacting protein (myomegalin) | WNA | Y | Hyperactivity, hemoglobin, bone mineral, retina/cornea |
| *Wdfy3* | 5:101980822:102217787 | WD repeat and FYVE domain containing 3 | SA | Y | Craniofacial/liver/spleen/brain/blood vessel morphology |
| *Fry* | 5:150042110:150421218 | FRY microtubule binding protein | WNA | Y | Mitosis |
| *Dpp6* | 5:27022355:27932498 | dipeptidylpeptidase 6 | WNA | Y | Abnormal nervous system morphology |
| *Akap9* | 5:3977410:4130204 | A kinase anchor protein 9 | SA | Y | Homeostasis, cholesterol, calcium, spermatogenesis, tooth morphology, body weight, immunity, fat |
| *Mical3* | 6:120908668:121107959 | microtubule associated monooxygenase, calponin and LIM domain containing 3 | ENA | Y | Immunity, activity |
| *Tspan11* | 6:127864585:127930940 | tetraspanin 11 | ENA | Y | Coat/hair, vocalization |
| *Met* | 6:17463799:17573979 | met proto-oncogene | WNA, SA | Y | Metabolism, cancer, pulmonary function/morphology, liver, brain |
| *Cdh8* | 8:99751103:100143103 | cadherin 8 | WNA | Y | Thermal nociception, tail movement, abnormal nervous system physiology |
| *Mlh1* | 9:111057296:111100854 | mutL homolog 1 | ENA, SA | Y | Fertility, immunity |
| *Wdr72* | 9:74017608:74190485 | WD repeat domain 72 | ENA | Y | Body weight, enamel/tooth |
| *Fat2* | 11:55141435:55227390 | FAT atypical cadherin 2 | ENA | N | Cell-cell adhesion |
| *Slc35b1* | 11:95275696:95282602 | solute carrier family 35, member B1 | WNA | N | Energy metabolism |
| *Tspan13* | 12:36064554:36092477 | tetraspanin 13 | WNA | Y | Bone density/structure, body fat, glucose |
| *Hivep1* | 13:42205304:42338504 | human immunodeficiency virus type I enhancer binding protein 1 | SA | Y | Heart morphology |
| *Epha3* | 16:63363897:63684538 | Eph receptor A3 | WNA | Y | Heart, spleen, body size |
| *Rorb* | 19:18907969:19088560 | RAR-related orphan receptor beta | ENA, WNA | Y | Eye, body size, coordination, regulation of circadian rhythm |

\*ENA: Phifer-Rixey *et al.* 2018; WNA: Ferris *et al.* 2021
\*\* Linked to differential expression, allele specific expression, and/or *cis*-QTL in comparisons among either wild mice from populations at different latitudes in the Americas (Mack *et al* 2018) or lab strains derived from populations at different latitudes in the Americas (Phifer-Rixey *et al.* 2018; Ballinger et al. 2023; Durkin *et al.* 2023)

### GWAS for body weight and body mass

Consistent with Bergmann’s rule, body size in house mice varies clinally in both North America and South America, with larger animals farther from the equator (Figure 5ab; S3; Phifer-Rixey *et al*. 2018; Suzuki *et al*. 2020; Ferris et al. 2021; Ballinger and Nachman 2022). Body weights of wild-caught mice are positively correlated with degrees from the equator in ENA and SA but not in WNA, as previously reported (Suzuki *et al*. 2020; Figure 5). However, lab-born descendants of mice from Edmonton are significantly larger than lab-born descendants of mice from Tucson (Ferris *et al*. 2021), consistent with patterns seen in ENA (Phifer-Rixey *et al*. 2018). To identify genes contributing to variation in body weight, we performed an association study using the exome data from 114 adult mice from South America (n=38), ENA (n=36), and WNA (n=40; Fig. S3). We excluded juvenile animals, pregnant or lactating females, and mice from Manaus who were housed in a laboratory setting before weighing. After filtering (MAF of 5% or greater and less than 10% missing data), 163,121 SNPs remained. We repeated the analysis using body mass index (BMI).

**Figure 5.**
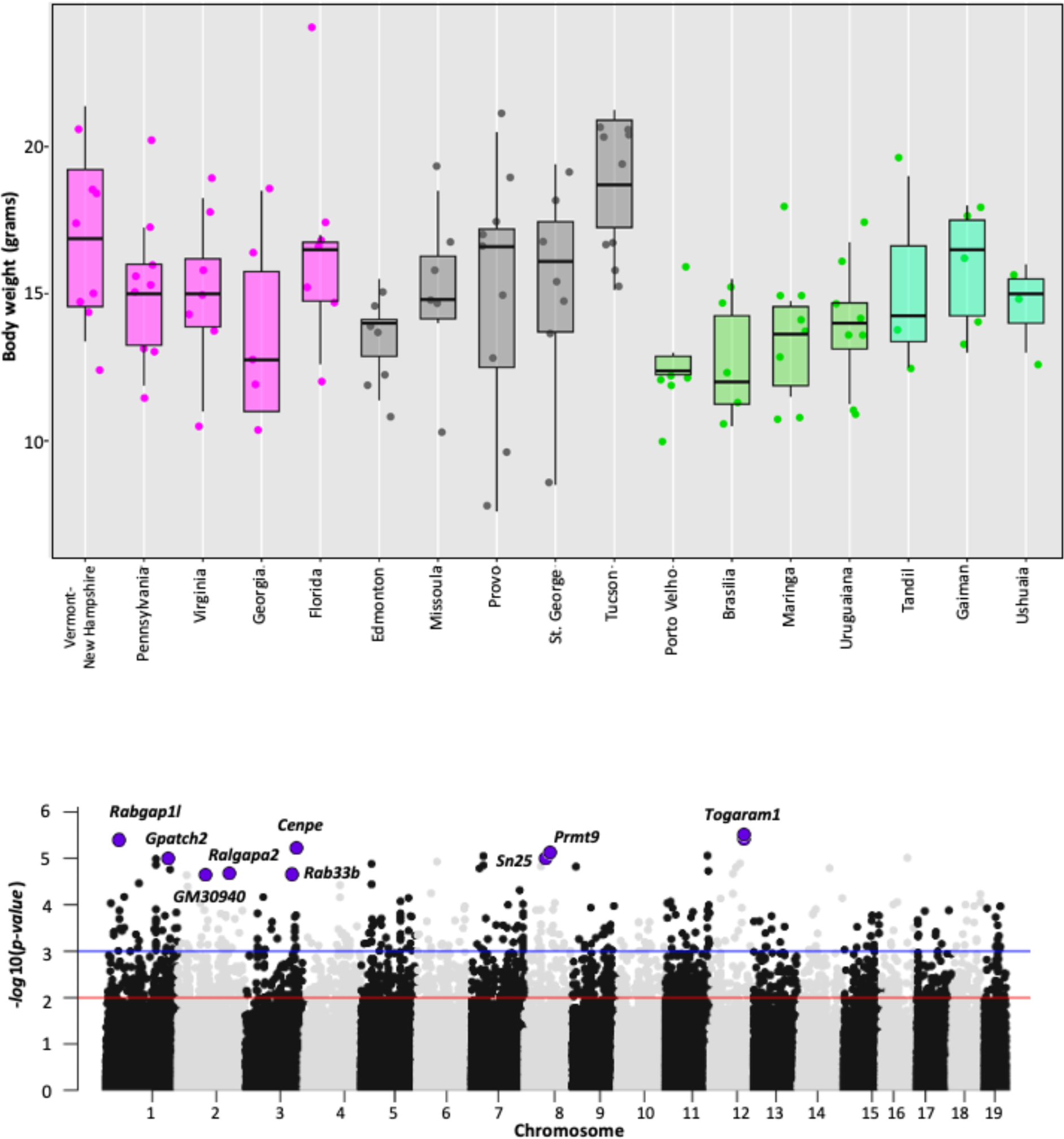
Body weight and genes associated with body weight in house mice from North and South America. **a)** Boxplot shows the distribution of adult body weights for each population from Eastern North America (magenta), Western North America (grey), and South America (green). Pregnant females have been excluded. **b)** Manhattan plot showing the genome-wide association results using exome and body weight data. Significant candidate SNPs (FDR ≤ 0.05) associated with body weight are highlighted in purple with the gene annotated for the SNP (red line indicates *p-value* = 0.01, and blue line indicates a *p*-*value* = 0.001). Description of the genes is in Table 3 and detailed GEMMA results are given in Table S13.

**Table 3.**
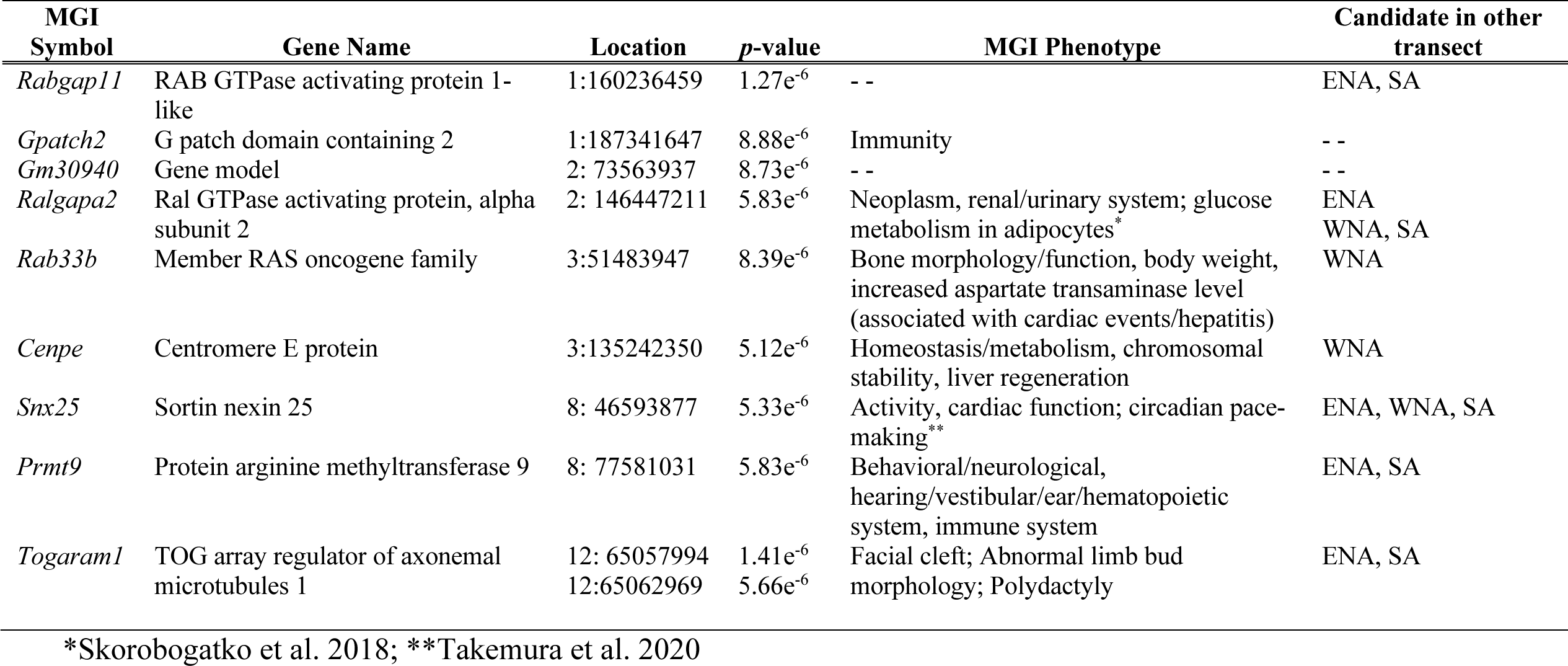
Genes annotated to candidate SNPs identified via GEMMA analysis of body weight across the Americas. Functional summarization is based primarily on MGI Mammalian Phenotype annotations and is not exhaustive. For additional detail on GEMMA results, see Table S13.

We carried out the analysis via GEMMA (Zhou and Stephens 2012) using sex as a covariate, controlling for relatedness among individuals testing, and correcting for multiple testing (false discovery rate of 5%). Using a false discovery rate correction, we did not find significant candidate SNPs associated with BMI. We found 10 SNPs in eight genes and one gene model (*Cenpe, Gm30940, Gpatch2, Rab33b, Rabgap1l, Ralgapa2, Prmt9, Snx25, Togaram1*) significantly associated with body weight (FDR ≤ 0.05, Tables 3, S13). None of these genes were identified in a similar GWAS limited to the western North American transect (Ferris *et al*. 2021), but all but one was identified as a candidate gene in scans for selection (Tables 3, S13). Overlap between the GWAS and selection scan candidates is significantly more than expected by chance (p < 0.005). Mutants in these genes relate to varied phenotypes, including aspects of metabolism, immunity, and morphology (MGI, Motenko *et al*. 2015). For example, mutations in *Cenpe* are associated with phenotypes related to lipase and glucose levels and *Cenpe* was differentially expressed in fat tissue collected from lab raised mice from New York and Florida (Phifer-Rixey *et al*.2018). Immunity phenotypes are annotated to *Gpatch2, Rab33b,* and *Prmt9.* Mutations in *Rab33b* also relate to bone morphology, body weight and cardiac function and *Prmt9* is also annotated to phenotypes relating to metabolism and grip strength (Table 3). *Snx9* and *Ralgapa2* were identified as candidates in all three transects and have annotated phenotypes relating to activity levels and tumor incidence, respectively. Mutations in *Togaram1* are associated with morphological phenotypes (e.g., facial cleft, abnormal limb bud morphology, and polydactyly; Table 3).

## Discussion

### Genetic variation in the Americas

We now have a comprehensive view of genomic variation in wild house mice across the Americas (Phifer-Rixey *et al*. 2018; Ferris *et al*. 2021; Fig. 1a, Fig. S4). Our data strongly support that the populations surveyed in Central and South America derive from *M. m. domesticus*, the subspecies found in Western Europe, consistent with known European colonization of the Americas (Boursot *et al*. 1993; Agwamba and Nachman 2023). Overall, contributions from other subspecies appear to be restricted to southwestern North America, with signals of admixture from *M. m. castaneus* in Tucson, AZ (Ferris *et al*. 2021). Previous studies of retroviral resistance also showed that mice in the Lake Casitas region of southern California showed significant introgression from *M. m. castaneus* (Orth *et al*. 1998) which may be linked to immigration from China (Gardner *et al*. 1991). Gene flow between southern California and Tucson would have been facilitated by the launch of a railroad line linking the areas in 1880 (Eubank 2016). Additional sampling in California and the Southwest would help determine the extent to which introgression from *M. m. castaneus* has contributed to genetic variation in Western North America.

We also found clear patterns of genetic differentiation among populations and among the three transects, with evidence of isolation by distance restricted to SA. Importantly, patterns of genetic variation across the continents suggest that the three sampled transects have independent recent evolutionary histories. Levels of differentiation among populations were not predicted by geographic distance in North America (Phifer-Rixey *et al*. 2018; Ferris *et al*. 2021). Idiosyncratic patterns may reflect complex but porous barriers to long distance gene flow due to human mediated transport (contemporary and/or historic). In contrast, geographic distance was a good predictor of genetic distance in South America, which may reflect more effective political and physical barriers (e.g., elevation, regional climates, water ways) to gene flow within the transect. Interestingly, Brazil and Argentina formed two clades sister to the Mexican clade. The clustering of Mexico with South American populations is somewhat surprising based on the significant physical barriers between Mexico and populations in Brazil and Argentina and the comparative proximity to southern populations in the WNA transect. While speculative, the existence of the South/Central American clade suggests the hypothesis that historic patterns of European colonization have influenced population structure in house mice in the Americas. These data complement genomic data from European populations, opening the door to a higher resolution understanding of the population genetics of an invasive species (Agwamba and Nachman 2023).

### Environmental adaptation in South America

In South America as in North America, we identified signals of selection associated with variation in latitude and mean annual temperature, with many of the candidate genes related to metabolism, fat, and body size, consistent with the observed variation in body size among mice from different latitudes (Suzuki *et al*. 2020; Figure S3). Top candidate genes were also linked to phenotypes related to immunity and cardiac, eye, and renal/urinary systems. We also considered PDM, identifying candidate genes with diverse functions including those related to immunity, muscle function, and kidney function/morphology, cholesterol, and sensory perception.

Nearly half of top candidate genes associated with variation in latitude and mean annual temperature in South America were also identified as candidates in previous studies of environmental adaptation in mice in the Americas and over half of them have been linked to differences in gene expression either in lab strains derived from these populations in the Americas or from wild populations in the ENA transect. Ten of these genes have mutant/knockout phenotypes related to blood/glucose/lipid homeostasis. These genes represent excellent candidates for additional investigation of the genetic basis of environmental adaptation in South America.

### Parallel adaptation across latitudinal gradients

With data from three transects across two continents, we can address how much of the response to selection across latitudinal gradients is shared and what that can tell us about adaptation in this system. Two major findings emerge. First, changes in regulatory regions dominate signals of selection in all three transects. While the relative contribution of regulatory and amino acid changing mutations to differences in phenotype is not known, the overwhelming and consistent signal suggests a key role for regulatory changes in rapid adaptation to climate across latitude. Second, while most candidate genes are unique to individual transects, there is compelling evidence for parallel adaptation in this system. Overlap among candidate genes from the three transects was significantly more than expected by chance for both latitude and mean annual temperature, with ∼16% of candidates shared between any two transects on average and ∼7% between all three. Adaptation across latitudinal gradients undoubtedly encompasses changes in many different traits, many of which are likely complex, such as body size, aspects of metabolism, immunity, and behavior. Parallel changes are expected to be more common for simple traits in which the mutational targets are small compared to complex and highly polygenic traits (Schluter *et al*. 2021). However, in a meta-analysis of published studies across a range of taxa, Conte *et al*. (2012) showed that the probability of parallelism increases with decreasing age of the common ancestor of the compared taxa. In this case, house mice in the Americas are of very recent origin, and the response to selection is almost certainly fueled by standing genetic variation from European populations.

### Connecting genotype to phenotype for adaptive complex traits

While body size, ear length, tail length, nesting behavior, and aspects of metabolism are known to vary with latitude across the Americas (Lynch 1992; Phifer-Rixey *et al*. 2018; Suzuki *et al*. 2020; Ballinger and Nachman 2022), genome-wide scans are agnostic to phenotype, allowing us to capture shifts driven by selection even when phenotypic variation may go unnoticed or be difficult to characterize. One challenge of genome scans is that it can be difficult to then connect candidate genes to specific functions, especially for complex traits for which effects of individual variants are expected to be small. Studies of wild house mice in the Americas have been integrative (e.g., Phifer-Rixey *et al*. 2018; Mack *et al*. 2018; Ferris *et al*. 2021; Ballinger *et al*. 2023; Durkin *et al*. 2023), bringing together multiple methods to help make connections between genetic and phenotypic variation for traits that impact fitness. The depth of functional work in house mice as a genetic model can help link candidates to function via mutants, knock-outs, and gene ontologies, for example (e.g., Motenko *et al*. 2015). These kinds of connections have helped generate hypotheses about genes and traits that contribute to environmental adaptation in this system (Phifer-Rixey *et al*. 2018; Ferris *et al*. 2021). For example, while many candidate genes relate to phenotypes known to vary with latitude, such as body size, the results of functional analyses suggest many additional phenotypes to consider, such as those relating to immunity, circadian rhythm, temperature sensing, and cardiac function. In addition, using gene expression as an intermediate phenotype can help identify specific candidate variants that are linked to differences in expression either in wild mice from different latitudes (Mack *et al*. 2018) or newly wild-derived laboratory strains from the Americas (Phifer-Rixey *et al*. 2018; Ballinger *et al*. 2023; Durkin *et al*. 2023; Dumont *et al*. 2023). While likely underpowered in most wild systems, genome-wide association studies are another approach that can help identify candidate loci for adaptive, complex traits. In this case, GWAS across the Americas for body weight identified eight candidate genes. All but one of these genes was also identified in selection scans, and two of them were identified in all three transects—*Snx25,* which is linked to activity, cardiac function, and more recently, circadian pace-making (Takemura *et al*. 2020) and *Ralgapa2*, which is linked to neoplasm and glucose homeostasis (Skorobogatko *et al*. 2018). Four of the others are associated with body size/metabolism/morphology and one with immunity. The significant overlap between the selection scan and GWAS results highlights these genes as candidates underlying adaptation. Finally, parallel shifts across independent latitudinal transects provide powerful evidence that a core set of genes with diverse functions contribute to environmental adaptation in the Americas.

Moreover, these approaches can bolster each other. For example, we can bring together the candidate genes identified for latitude and MAT in all three transects with published functional data and gene expression studies (Phifer-Rixey *et al*. 2018; Mack *et al*. 2018; Ballinger *et al*. 2023; Durkin *et al*. 2023). *Mlh1* was a candidate in all three transects and a top candidate in two. Mutants in *Mlh1* affect aspects of immune system function among other phenotypes. Laboratory studies of inbred strains from the Americas point to expression differences in *Mlh1* among strains (Ballinger *et al*. 2023; Durkin *et al*. 2023) and wild mice from opposite ends of the ENA transect (Mack *et al*. 2018) and it was associated with a *cis*-eQTL in ENA. More broadly, gene ontology enrichment analyses point to a role for cardiovascular, immunity, and retinal traits in adaptation and candidates like *Mc3r*, *Mgat4*, *Akap9*, and *Rorb*, point to a role for metabolism, body size, and circadian rhythm.

Multiple studies (Mack *et al*. 2018; Ballinger *et al*. 2023) have supported a role for *Bcat2* and *Adam17* in adaptive divergence in body size in this system. Mutants for both genes affect body mass (Wu *et al*. 2004; She *et al*. 2007; Gelling *et al*. 2008; Blake *et al*. 2017) and both were identified as candidates associated with latitude and linked via patterns of genetic variation and gene expression to variation in body size in the ENA transect (Mack *et al*. 2018). In South America, *Bcat2* was identified as a candidate gene for latitude and MAT. Interestingly, while *Adam17* was only identified as associated with latitude in selection scans in ENA, it was also identified as a candidate for PDM in South America. Mack *et al*. (2018) reported support for an important role for *Adam17* in body weight divergence among populations of house mice from eastern North America, and Ballinger *et al*. (2022) identified *cis*-regulatory changes at *Adam17* in comparisons between mice from Brazil and mice from New York (which also differ in body size).

Another example of the potential of this kind of integrative approach comes from the *Trpm* gene family. TRPM channels act as cellular sensors with impacts on diverse physiological processes, including temperature sensing, mineral homeostasis, cardiac rhythm, and immunity (Chubanov *et al*. 2023). Multiple genes in the *Trpm* family have been linked to response to environmental factors like temperature and light, among others (e.g., Vriens *et al* 2011; Held *et al*. 2015; Tan and McNaughton 2016; Yang et al. 2020). *Trpm2* was identified as a candidate gene for latitude and MAT in all transects (Phifer-Rixey et al. 2018; Ferris *et al*. 2021) and was linked to differences in gene expression in strains derived from different populations of the Americas (Phifer-Rixey *et al*. 2018). Mutant phenotypes in *Trpm2* relate to immunity (Zou *et al*. 2013) and experimental results link *Trpm2* to sensitivity to warmth (Tan and McNaughton 2016) and insulin secretion (Togashi *et al*. 2006). *Trpm6* was identified as a candidate in all three transects, functions in Mg+ transport, and was linked to differential expression under different temperature regimes in lab strains derived from New York and Brazil (Ballinger *et al*. 2023). *Trpm8* was identified as a candidate in both SA and WNA and has been shown to drive cold sensitivity in vertebrates (Matos-Cruz *et al*. 2017; Key *et al*. 2018; Yang *et al*. 2020). Finally, *Trpm3* and *Trpm1* were identified for the first time as candidates relating to latitude and MAT in the SA transect. *Trpm1* is linked to light reception (e.g., Morgans *et al*. 2008) and *Trpm3* has been shown to function in heat sensitivity (Vriens et al 2011; Held *et al*. 2015). *Trpm3* was differentially expressed in laboratory studies of inbred strains derived from New York and Brazil under different thermal conditions (Ballinger *et al*. 2023). Together, these results underscore the role of genes that mediate response to the environment in adaptation to novel climates.

## Conclusion

House mice arrived in the Americas in association with human colonization, quickly and successfully establishing populations in a variety of climates and habitats. Perhaps as expected given their niche as human commensals, patterns of genetic variation across the Americas suggest major impacts of human migration on population structure. Also as predicted by their natural history, genetic differentiation among populations is relatively high and, while evidence for isolation by distance is restricted to South America, populations across the Americas cluster by continent and by region within continent. Bringing together results from three independent transects, we demonstrate that adaptation across latitudinal gradients in the Americas is largely driven by unique changes and largely in regulatory regions, but significant overlap among transects provides evidence for parallel adaptation and highlights a core group of candidate genes. The wealth of functional data available for house mice together with gene expression studies in wild populations and new wild-derived inbred strains from the Americas help connect these candidates to traits with potential to impact fitness. While much more is now known about the genetics of wild house mice in the Americas, population genomic data at this geographic scale combined with the functional resources generated (new wild-derived strains, phenotype data, and gene expression data) point to great potential for continued investigation of the genetic basis of adaptation in this system and more broadly, the connection between genotype and phenotype in house mice (Dumont *et al. 2023*).

## Materials and Methods

### Sampling

Mice were collected using live Sherman traps from sites within eight sampling locations along a latitudinal transect in South America and from two sampling locations in Mexico (Figure 1a; Table S1). When possible, mice were collected from ten or more sites within each sampling location and sites were at least 500 m apart to avoid the inclusion of close relatives. In some sampling locations, this collection scheme was not tractable and either fewer sites were included (Ushuaia, Argentina; Chiapas, Mexico) or some sites were less than 500 m apart (Table S1). Sex and body size data were recorded for each mouse along with latitude, longitude, and elevation (Table S1). Measures of size included total length, tail length, hindfoot length, and ear length as measured with a ruler and total weight (grams) measured using a micro-line spring scale. Animals were sacrificed in accordance with a protocol approved by the Institutional Animal Care and Use Committee (IACUC) of the University of California, Berkeley. Tissues including the liver, kidneys, and spleen were collected and either stored in liquid nitrogen or dry ice until transfer to a -80°C freezer or immersed in 96% EtOH that was drained and replaced after 24 hrs and then stored at 4°C. Skins, skulls, and skeletons were deposited in the Museum of Vertebrate Zoology, University of California, Berkeley (Table S1).

### DNA Extraction, library preparation, and sequencing

DNA was extracted and exome capture libraries were prepared as in Phifer-Rixey et al. (2018). Individuals were pooled for capture and each pool of enriched capture libraries was then sequenced on each of five lanes of an Illumina HiSeq4000 (150-bp paired-end) resulting in an average of approximately 6.4 GB of raw data per individual (Table S1). Sequence data from three individuals (FMM273, FMM 275, FMM276, Rodonia, Brazil) was generated separately but with a similar capture, pooling, and sequencing approach.

### Exome-capture pipeline

The exome sequence data were cleaned, and adapters were removed with the program AdapterRemoval (Schubert *et al*. 2016) using a minimum quality of PHRED ≥ 30. Then, we used the *Escherichia coli* genome (ASM584v2) to filter potential contamination. We mapped the exome raw reads against the *E. coli* genome, and we retained the unmapped reads using HISAT2 (Kim *et al*. 2019). After cleaning and filtering, we retained more than 99% of the reads (Table S1). On average the sequence depth coverage per site was 33.6x and 92% of the targeted exome was covered. The resulting reads were mapped to the house mouse reference genome (GRCm38.p6) using BWA-MEM (Li and Durbin, 2010; Jo and Koh 2015). Aligned reads were sorted, duplicates were marked and removed, and the reads that aligned to chromosomes X and Y were extracted using Picard and Samtools software (https://broadinstitute.github.io/picard/) (Danecek *et al*. 2021). We followed the GATK Best Practices pipeline to identify artifacts or technical errors made by the sequencing machine using base calibrator tools (BSQR and ApplyBQSR). Additionally, we performed a local realignment for indels and a variant filtering using the variant quality score recalibrations (VSQR) (Van *et al*. 2013).

Based on the Bayesian method implemented in ANGSD (Korneliussen *et al*. 2014; Durvasula et al., 2016), we estimated the allele frequencies and called SNPs for the six populations. We used the recalibrated bam files, a posterior probability of 95%, a p-value of the likelihood ratio test of 1x 10^-6^ (SNP_pval), and we removed those sites with a minimum allele frequency (minMaf) less than 5%. Finally, we retained those variants that were present in at least 80% of the samples, obtaining a total of 271,720 SNPs.

### *Mus sp.* Genomic data and admixture analysis

To investigate genetic admixture between South American and North American populations and the historical relationships among *Mus musculus*, we included the previously published genomic information for 50 individuals of *M. m. domesticus* from Eastern North America (Phifer-Rixey *et al*. 2018), 50 individuals from Western North America (Ferris *et al*. 2021), 10 *M. m. domesticus* from France and Germany, three *M. m. musculus* individuals each from the Czech Republic and Kazakhastan, and 10 individuals of *M. m. castaneus* (Harr *et al*. 2016; Table S2). All of these data are publicly available and were downloaded from the Sequence Read Achieve (SRA), and the European Nucleotide Archive (EnA) repositories (Table S2).

We cleaned and filtered the raw reads using the same method applied to the data generated for this study. The resulting reads were mapped to the house mouse reference genome (GRCm38.p6) using BWA-MEM. The mapped reads were sorted, duplicates were marked and removed using picard and samtools software. We used GATK pipeline to identify artifacts or technical errors (BSQR and ApplyBQSR). Additionally, we performed a local realignment for indels and a variant filtering using the variant quality score recalibrations (VSQR) (Van *et al*. 2013). Finally, from the genomic bam files, we extracted the exome coordinate regions using samtools, bedtools and bash scripts (Li *et al*. 2009; Quinlan and Hall, 2010).

We used the autosomal recalibrated bam files and the software ANGSD (Korneliussen *et al*. 2014) to calculate genotype likelihoods for polymorphic sites, applying a SNP_pval 2e-6 and minMaf of 5% minMaf. To obtain an accurate estimate of the admixture proportions and the best genetic cluster value (K), we ran *NGSadmix* for several K values: K=2 to K=5, with 5,000 as the maximum number of EM iterations. We used the log-likelihood estimated for each K to calculate the Cluster Markov Packager Across K from Evanno using R scripts.

### Phylogenetic reconstruction

To investigate the phylogenetic relationships between the South American house mice populations and their close relatives from North America, we used the exome sequence data generated for this study, 86 individuals from Mexico and South America, and the exome data from the 100 individuals from North America included in the admixture analysis (Phifer-Rixey *et al*. 2018; Ferris *et al*. 2021). We also incorporated the genomic data of *M. spretus* (one individual, Project: PRJEB11742, sample: ERR1124353) (Table S2), as an outgroup.

We used Haplotype Caller from GATK pipeline to perform the SNP calling in autosomal genomic regions for all the individuals of *M. m. domesticus* and *M. spretus.* We carried out the SNP calling first using all the individuals and then excluding Mexico due to small sample size. We filtered to only include the most suitable SNPs for phylogeny construction, using Variant quality score recalibration (VQSR) and QD>= 20 (QualbyDepth) to identify variants. Only biallelic sites were retained. Additionally, we pruned SNPs for linkage disequilibrium using non-overlapping 50 kb sliding windows, and an r^2^ = 0.5 (50 50 0.5), allowing a missing data cut-off of 20%. Recalibrated bam files, ngsDist from *NGS* Tools (Korneliussen *et al*. 2014), and RaxML (Stamatakis 2014) were used to build the phylogenetic tree, given *M. spretus* as the outgroup and 100 repetitions of bootstrapping for node support. FigTree software (Rambaut 2009) was used to visualize the tree.

### Genetic differentiation between populations

To explore population structure in the house mouse populations from North America, Mexico and South America, we used ngsCovar software (Fumagalli *et al*. 2014) to generate a genetic covariance matrix from the genotype posterior probabilities (using a snp *p-value* = 1×10^-3^ and 5% minimum allele frequency) generated from the autosomal bam files of 186 individuals. We used R to perform the eigenvalue decomposition and generated the PCA plot using the three principal components. With ANGSD, we estimated *F_st_* for each pair of populations (North and South America), using the unfolded pairwise site frequency spectra (SFS) as priors for the allele frequency probabilities at each site. Additionally, we used VCFtools to calculate pairwise Weir and Cockerman’s *F_st_* for all the populations. Finally, for the populations from South America, we analyzed the relationship between genetic distance and the geographic distance by performing a Mantel Test using the R packages vegan, adegenet and hierfstat (Jombart 2008; Goudet 2015).

### Relatedness analysis

To infer relatedness between pairs of individuals in the South American populations, we used ANGSD to estimate the genotype likelihoods and allele frequencies (snp p-value 1×10^-6^, and 5% of minor frequency allele) and the program ngsRelate (Hanghøj *et al*. 2019) to calculate different relatedness and inbreeding coefficients. To identify close relatives, we used the relatedness coefficient proposed by Hedrick and Lacy (2015). Those pairs of individuals with a relatedness measure above 0.25 were considered relatives (Fig. S3). Relatedness heatmaps were generated using R (R Team Core).

### Environmental association analysis

We used the Arctos database to obtain the geographic coordinates of each locality (latitude and longitude) for the 186 individuals of house mice from South America, eastern and western of North America (Table S1). Based on the geographic coordinates, we extracted the 19 bioclimatic variables from WorldClim database (30 seconds spatial resolution; Fick and Hijmans, 2017) for each individual, using the package raster in R (Bivand, 2011). We performed a PCA using the 19 bioclim variables to explore the climatic variability of our samples and to identify the most informative environmental variables (Table S5). Moreover, we tested the relationship between latitude and the two bioclim variables (Bio 1-Annual Mean Temperature and Bio 14 – Precipitation of the Driest Month) calculating Pearson’s correlations.

To identify candidate genes underlying environmental adaptation, we performed a population genomic scan for selection using the Latent Factor Mixed Model (LFMM) software (Frichot *et al*. 2013) and three environmental variables (Latitude, Bio 1, and Bio14). LFMM used a Bayesian bootstrap algorithm that accounts for population structure while identifying genetic polymorphisms that exhibit correlations with variables of interest (environmental measures or phenotypic traits). For each environmental variable, we ran LFMM (25 repetitions, burn-in = 100,000 and 500,000 iterations), using K=3 for South America, and a K=2 for the eastern and western transect. For each LFMM result, we estimated the genomic inflation factor (λ), and the *p-values* were adjusted to control for the false discovery rate (FDR). We identified candidate SNPs using a threshold *q-value* ≤ 0.05 and *z-score* ≥ 2.

To explore the potential functional significance of the candidate SNPs identified, we used Ensembl’s Variant Effect Predictor command line software (McLaren *et al*. 2016) with the house mouse reference genome (GRCm38.p6). Many SNPs had more than one potential functional consequence. To classify the SNPs annotated based on their “primary” functional consequence we followed the scheme proposed by Phifer-Rixey et al., (2018): 1) missense, stop lost or stop gain; 2) 3’ or 5’ UTR; 3) synonymous; 4) non-coding exon variants or non-coding transcript variants; 5) intron or splice site variants; 6) downstream or upstream variants.

Additionally, we extracted from The Mouse Genome Informatics Database (MGI; Blake *et al*. 2021) the gene ontology terms (GO) (Hayamizu *et al*. 2015) associated with the genes annotated in each LFMM analysis. Using the R package topGO (Alexa and Rahnenfuhrer 2023), we performed an enrichment analysis, and the *p-values* were adjusted using the Fisher’s exact test and the false discovery rate. We generated a network correlation of the most significant nodes obtained with the package topGO and networkplot in R.

We used the R package “VennDiagram” (Chen and Boutros 2011) to identify those genes shared among LFMM environmental variables results, and also among transects. With the R package “RegionR” (Gel *et al*. 2016), we performed permutation analysis to test if the genes shared between transects (South America – ENA, South America – WNA, ENA – WNA) were significantly greater than expected by chance, running 100,000 replicates with replacement.

We estimated the allele frequencies for the eight populations from South America (Manaus, Porto Velho, Brasilia, Maringa, Uruguaiana and Ushuaia; 76 individuals) using the samtools. Based on the list of candidate SNPs (*q-value* ≤0.05 and *z-score* ≥ 2), we performed linear regressions between the allele frequencies for each candidate SNPs and latitude, and we retained those SNPs with a *R^2^* ≥ 0.65 and *p-value* ≤ 0.01.

### Genome-wide association analysis for body weight

We conducted a genome wide association analysis to identify genes that contribute to variation in body weight using the program GEMMA (Zhou and Stephens, 2012) that implements an algorithm of linear mixed models. We used the body weight data from 116 adult house mice from the Americas. We excluded individuals reported as juveniles, subadults, pregnant, those with undetermined reproductive status, and those that were weighed after being housed in a laboratory colony. We used a total of 163,121 loci (filtered by maf > 5%, and < 10% of missing data) to generate a genotype matrix as an input for GEMMA. We ran GEMMA using the body weight as a phenotype, controlling for relatedness among individuals and population structure, and using sex as a covariate to reduce the number of false positives. We adjusted *p-values* using the false discovery rate method. Finally, the SNP candidates were annotated using the Ensemble’s variant predictor command line program. To test whether the overlap between GWAS candidates and the selection scans was more than expected by chance, we used a permutation test implemented in RegionR (Gel *et al*. 2016) as described above.

## Data availability

Exome reads sequencing are available at the Sequence Read Archive (SRA) from NCBI, BioProject accession PRJNA776897. Supplementary Table S1 includes the complete metadata of the samples: locality information, sex, reproductive state, individual body size measures, read sequencing information, and the Museum of Vertebrate Zoology vouchered specimen number and accession. The code and scripts used for the analysis are available from Github (https://github.com/YocelynG/HouseMouse_EnvAdapt).

## Acknowledgments

We thank the members of the Nachman Lab for their valuable comments and discussions. We thank Felipe Martins for extensive field work. We thank Katya Mack, Sylvia Durkin, and Mallory Ballinger for sharing work in progress. We thank Ticul Álvarez for providing samples from Mexico, Andreas Chavez and Taichi A. Suzuki for help with data collection, Libby Beckman for comment and discussion, and Lydia Smith and Ke Bi for technical support and expertise. This work was facilitated by access to the Patung cluster, LANCIS-Instituto de Ecología, UNAM.

## Supporting information

**Alternative language abstract**

Resumen (Spanish)

Resumo (Portuguese)

**Alternative language author summary**

Resumen del autor (Spanish)

Resumo do autor (Portuguese)

## Supplementary Tables

**Supplementary Table 1**. List of 86 wild-caught house mice individuals (*Mus musculus domesticus*) collected in Mexico (N=10), Brazil (N=60) and Argentina (N=26). Table contains: Collector’s number, SRR ID, Museum of Vertebrate Zoology catalog number, exact collecting locality, latitude, longitude, sex, reproductive data, measurements of length and body weight, and data of exome sequencing (number of reads, length of reads and coverage)

**Supplementary Table 2**. Sample information for the European samples of *Mus musculus musculus, Mus musculus domesticus, Mus musculus castaneus,* and *Mus spretus* from Harr et al. (2016; Doi: 10.1038/sdata.2016.75) included in our analyses.

**Supplementary Table 3**. Pairwise differentiation (Fst) across the three transects: South America, East and West of North America.

**Supplementary Table 4**. Values for bioclimatic environmental variables for sampled populations of *M. musculus domesticus* across the Americas.

**Supplementary Table 5**. Loadings of bioclimatic variables for the first five principal components for the analysis of climate data for all included populations in the Americas.

**Supplementary Table 6**. Results of Latent Factor Mixed Model (LFMM) analysis for the each of three variables (latitude, MAT, PDM) in South America populations as well as information about top candidates and shared candidates, and the allele frequencies.

**Supplementary Table 7**. The distribution of candidate SNPs identified in LFMM analyses of South American populations and all SNPs included in the analyses across predicted functional consequence category.

**Supplementary Table 8**. The results of enrichment analysis for candidates identified in LFMM analysis of South American populations.

**Supplementary Table 9**. Classification and annotation of SNPs identified as candidates in LFMM analyses of North American populations with LAT, MAT, and PDM.

**Supplementary Table 10**. Proportion of genes shared between and across all transects for each variable.

**Supplementary Table 11**. Pairwise permutation test results for overlap among candidate genes for each environmental variable identified across the three transects: South America (SA), Eastern of North America (ENA), and Western of North America (WNA), using a *p-value* £ 0.05 and 10,000 permutations. The number of genes shared for each variable are described in Figure 4.

**Supplementary Table 12**. Functional information for candidate genes shared across the three transects for each environmental variable.

**Supplementary Table 13**. Gene annotations and functional information for candidate SNPs identified via GEMMA for body weight.

**Supplementary Figure 1. a)** Climatic variation across the sampled localities in South America (SA), East (ENA) and West (WNA) of North America using PCA with 19 bioclimatic variables from the WorldClim database. The first component is mainly associated with variation variables relating to temperature (e.g., mean annual temperature, MAT), and the second principal component was mainly associated with precipitation of the driest month (PDM) and precipitation of the driest quarter. **b)** Latitude and Bio1-MAT are significantly correlated across the sampled localities (ENA, WNA, SA). There is no evidence of correlation between **c)** latitude and Bio14-PDM and **d)** MAT and Bio14-PDM across the sampled localities (ENA, WNA, SA).

**Supplementary Figure 2**. Heatmap of pairwise relatedness coefficients between individuals from the same population using the relatedness estimator R_AB_, described by Hedrick and Lacy (2014). Individuals that were removed because they were close relatives to another sampled mouse (with a pairwise relatedness value greater than 0.25) are shown in bold.

**Supplementary Figure 3**. Body weights of adult mice across Eastern of North America (ENA), Western of North America (WNA), and South America (SA) included in the GEMMA analysis.

**Supplementary Figure 4**. Structure plot showing population genetic clusters using K=2:7 across North and South America.

**Supplementary Material**. LFMM output, zscores, p-value correction, and allele frequencies for South America, Eastern and Western of North America transects.

## Supporting Information

Alternative language abstract (Spanish and Portuguese)

Alternative language author summary (Spanish and Portuguese)

